# Nucleus Specific expression in the multinucleated mushroom-forming fungus *Agaricus bisporus* reveals different nuclear regulatory programs

**DOI:** 10.1101/141689

**Authors:** Thies Gehrmann, Jordi F. Pelkmans, Robin A. Ohm, Aurin M. Vos, Anton S. M. Sonnenberg, Johan J. P. Baars, Han A. B. Wösten, Marcel J. T. Reinders, Thomas Abeel

**Author notes:** Current position: CBS/KNAW Westerdijk Fungal Biodiversity Institute, The Netherlands. Current position: Department of Biotechnology, Delft University of Technology, The Netherlands (Aurin M. Vos).

## Abstract

**Motivation:** Fungi are essential in nutrient recycling in nature. They also form symbiotic, commensal, parasitic and pathogenic interactions with other organisms including plants, animals and humans. Many fungi are polykaryotic, containing multiple nuclei per cell. In the case of heterokaryons, there are even different nuclear types within a cell. It is unknown what the different nuclear types contribute in terms of mRNA expression levels in fungal heterokaryons. Each cell of the cultivated, mushroom forming basidiomycete *Agaricus bisporus* contains 2 to 25 nuclei of two nuclear types, *P1* or *P2,* that originate from two parental strains. Using RNA-Seq data, we wish to assess the differential mRNA contribution of individual nuclear types in heterokaryotic cells and its functional impact.

**Results:** We studied differential expression between genes of the two nuclear types throughout mushroom development of *A. bisporus* in various tissue types. The two nuclear types, produced specific mRNA profiles which changed through development of the mushroom. The differential regulation occurred at a gene and multi-gene locus level, rather than the chromosomal or nuclear level. Although the P1 nuclear type dominates the mRNA production throughout development, the P2 type showed more differentially upregulated genes in important functional groups including genes involved in metabolism and genes encoding secreted proteins. Out of 5,090 karyolelle pairs, i.e. genes with different alleles in the two nuclear types, 411 were differentially expressed, of which 246 were up-regulated by the P2 type. In the vegetative mycelium, the P2 nucleus up-regulated almost three-fold more metabolic genes and cazymes than P1, suggesting phenotypic differences in growth. A total of 10% of the differential karyollele expression is associated with differential methylation states, indicating that epigenetic mechanisms may be partly responsible for nuclear specific expression.

**Conclusion:** We have identified widespread transcriptomic variation between the two nuclear types of *A. bisporus*. Our novel method enables studying karyollelle specific expression which likely influences the phenotype of a fungus in a polykaryotic stage. This is thus relevant for the performance of these fungi as a crop and for improving this species for breeding. Our findings could have a wider impact to better understand fungi as pathogens. This work provides the first insight into the transcriptomic variation introduced by genomic nuclear separation.

## Introduction

Fungi are vital to many ecosystems, contributing to soil health, plant growth, and nutrient recycling^1^. They are key players in the degradation of plant waste ^2,3^, form mutually beneficial relationships with plants by sharing minerals in exchange for carbon sources^4,5^ and by inhibiting the growth of root pathogens^6,7^. They even form networks between plants, which can signal each other when attacked by parasites^8^. Yet, some are plant pathogens responsible for huge economic losses in crops^9–11^.

The genome organization of fungi is incredibly diverse and can change during the life cycle. For instance, sexual spores can be haploid with one or more nuclei or can be diploid. Sexual spores of mushroom forming fungi are mostly haploid and they form monokaryotic (one haploid nucleus per cell) or homokaryotic (two or more copies of genetically identical haploid nuclei) mycelia upon germination. Mating between two such mycelia results in a fertile dikaryon (one copy of the parental nuclei per cell) or heterokaryon (two or more copies of each parental nuclei) when they have different mating loci^12^. In contrast to eukaryotes of other kingdoms, the nuclei do not fuse into di- or polyploid nuclei but remain side by side during the main part of the life cycle. Only just before spores are formed in mushrooms, do these nuclei fuse, starting the cycle anew.

*Agaricus bisporus* is the most widely produced and consumed edible mushroom in the world^2^. Heterokaryotic mycelia of the button mushroom *Agaricus bisporus* var. *bisporus* (Sylvan A15 strains) have between 2 and 25 nuclei per cell^13,14^ (Figure 1a-d). The genomes of both ancestral homokaryons have been sequenced^1,15^ showing that DNA sequence variation is associated with different vegetative growth capabilities^1^. Due to the two nuclear types, each gene exists at two alleles separated by nuclear membranes, which we call karyolleles. Although there have been a few studies investigating the expression of genetic variety in the transcriptome^16,17^, the differential transcriptomic activity of two (or more) nuclear types has never been systematically investigated in a heterokaryon at the genome wide scale. Based on SNPs identified in mRNA sequencing, it has been suggested that allele specific expression is tightly linked to the ratio of the nuclear types in a basidiomycete^18^.

**Figure 1.**
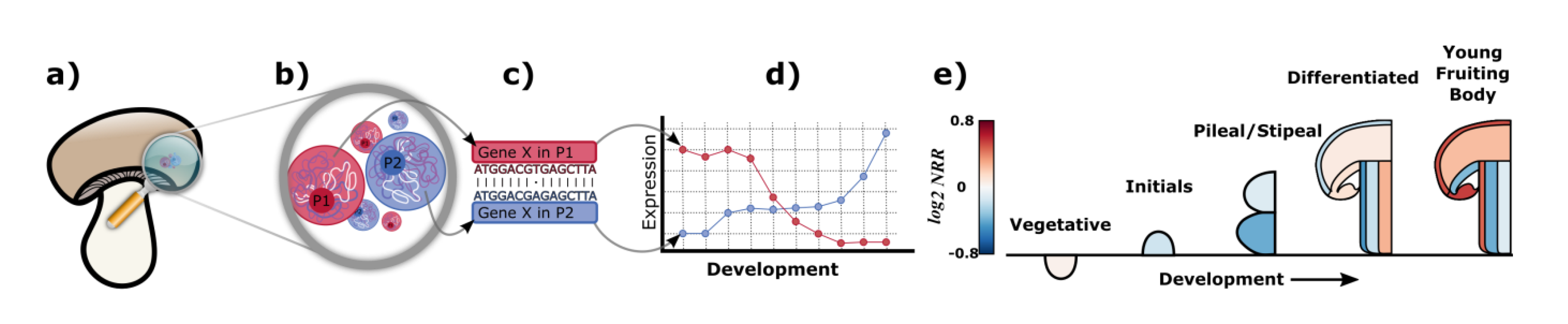
Nuclear type specific expression in A. bisporus. a) The A. bisporus mushroom is composed of different tissues that consist of hyphae comprised of cellular compartments. b) Each cellular compartment is a heterokaryon containing between 2 and 25 nuclei. In our strain, each nucleus is either of type P1 (red) or P2 (blue). Both nuclear types are haploid, and contain exactly one copy of each gene. However, because there are multiple nuclei, there may be multiple copies of each gene in the cell. c) Furthermore, the gene in the two types, which we call karyolleles, may differ in their genetic sequences. d) These differences in transcript sequence allow us to quantify expression of each karyollele in each tissue and to investigate nucleus specific expression. e) Read count ratios at the nuclear type level (Equation 5) of Agaricus bisporus throughout its development. Red colour indicates higher P1 activity, blue colour higher P2 activity. The scale bar indicates the log2 fold change in activity between the P1 and P2 nuclear types. We observe a differential mRNA activity in different mushroom tissues.

Allele specific expression in mononuclear cells has been studied in fungi^19^, plants^20^, animals^21^,and humans^22^. Such studies have shown that allele heterogeneity is linked to differential allele expression and cis-regulatory effects^21–23^, and even sub-genome dominance^24^.*A. bisporus* is in many ways an excellent model organism to investigate differential karyollele expression. It only has two nuclear types in the heterokaryon contrasting to the mycorrhizae that can have more nuclear types^25,26^, making computational deconvolution of mRNA sequence data intractable with currently available tools. Additionally, the recently published genomes of the two nuclear types of *Sylvan A15*^15^ exhibit a SNP density of 1 in 98 bp allowing differentiation of transcripts in high throughput sequencing data. Finally, bulk RNA-Seq datasets of different stages of development and of different tissues of the fruiting bodies are available^2, 27^

Here, we show that differential karyollele expression exists in *Agaricus bisporus Sylvan A15* strain,which changes across tissue type and development and affects different functional groups. Further, we show that differential karyollele expression associates with differential methylation states, suggesting that epigenetic factors may be a cause for the differential regulation of karyolleles.

## Results

### Karyollele specific expression through sequence differences

To assign expression levels to individual karyolleles, we exploit sequence differences between karyollele pairs in the P1 and P2 homokaryon genomes of *A. bisporus A15* strain (Materials). Briefly, the sequence differences define marker sequences for which the RNA-Seq reads uniquely match to either the P1 or the P2 variant, effectively deconvolving the mRNA expression from the two nuclear types (see Methods). There are a total of 5,090 distinguishable karyollele pairs between the *P1* and *P2* genomes, corresponding to ∼46% of all genes. The remaining genes could not be unambiguously matched, or the karyollele pairs had too few sequences differences. Most (80%) distinguishable karyollele pairs had the same number of markers in each homokaryon. For the remaining pairs (20%), the number of markers per karyollele was different (see Supplementary Material Note A). This variation can be explained by the non-symmetric number of markers produced by the different kinds of variation. While a SNP will result in one marker in each karyollele, an indel (if longer than 21bp) will result in one marker in one karyollele, and at least two in the other. Karyollele specific expression is expressed as a read count ratio that reflects the relative abundance of mRNAs originating from the P1 or P2 nuclear types (Equation 3, Methods).

We studied *A. bisporus*’ karyollele specific expression for different tissues and development in two RNA-Seq datasets, one studying the mycelium in compost throughout mushroom harvest, and one studying different mushroom tissues throughout mushroom formation (Figure 1e, Supplementary Material Notes B, and Materials). Measured difference in expression between nuclear types is not correlated with the number of markers (p > 0.05) for any of the samples, nor is it correlated with CG content (see Supplementary Material Notes C).

### P1 and P2 mRNA production differs per tissue and across development

First, we assess the total mRNA production of the P1 and P2 nuclear types and their relative contributions during development. To do this, we considered the total number of reads uniquely matching to P1 with respect to P2. Figure 1e shows that this nuclear type read count ratio (NRR, see Equation 5, Methods) changes throughout development and across tissue types. For example, during the *‘Differentiated’* stage, the P2 nuclei are dominant in the skin, but in the *‘Young Fruiting Body’*, the P1 nuclei dominate the skin (two right most panels in Figure 1e). In contrast, the *‘Stipe Center’* is dominated by P1 nuclei in the differentiated stage, while later the expression of P2 nuclei dominates.

The transcription patterns throughout the mushroom development differ between the karyolleles. Based on a principal component analysis of the expression profiles of each nuclear type, we observe that the expression profiles of P1 and P2 group together in different clusters, based on the first and second principal components (Supplementary Material Note D). This clustering is indicative of distinct regulatory programs. It appears as though the first principal component represents the tissue type, and the second represents the nuclear type. Interestingly, measurements of the same tissue from P1 and P2 do not have exactly the same value for the first principal component, indicating that the difference in nuclear type does not entirely explain the variation between P1 and P2.

### Within a sample, mRNA production of P1 and P2 vary between chromosomes

Figure 2a shows the Chromosome Read count Ratios (CRR, Equation 4), demonstrating that some chromosomes are more active in P1 (e.g. chromosome 8) throughout development, while others are more active in P2 (e.g. chromosome 9). Expression of other chromosomes depend on the developmental state, changing in time (e.g. chromosome 2). The chromosome log2 fold changes lie between [-0.60, 0.79]. In the vegetative mycelium we see less drastic differences in mRNA production throughout development than in the mushroom tissues, with expression log2 fold changes between [-0.28, 0.36] (see Supplementary Material Notes B).

**Figure 2.**
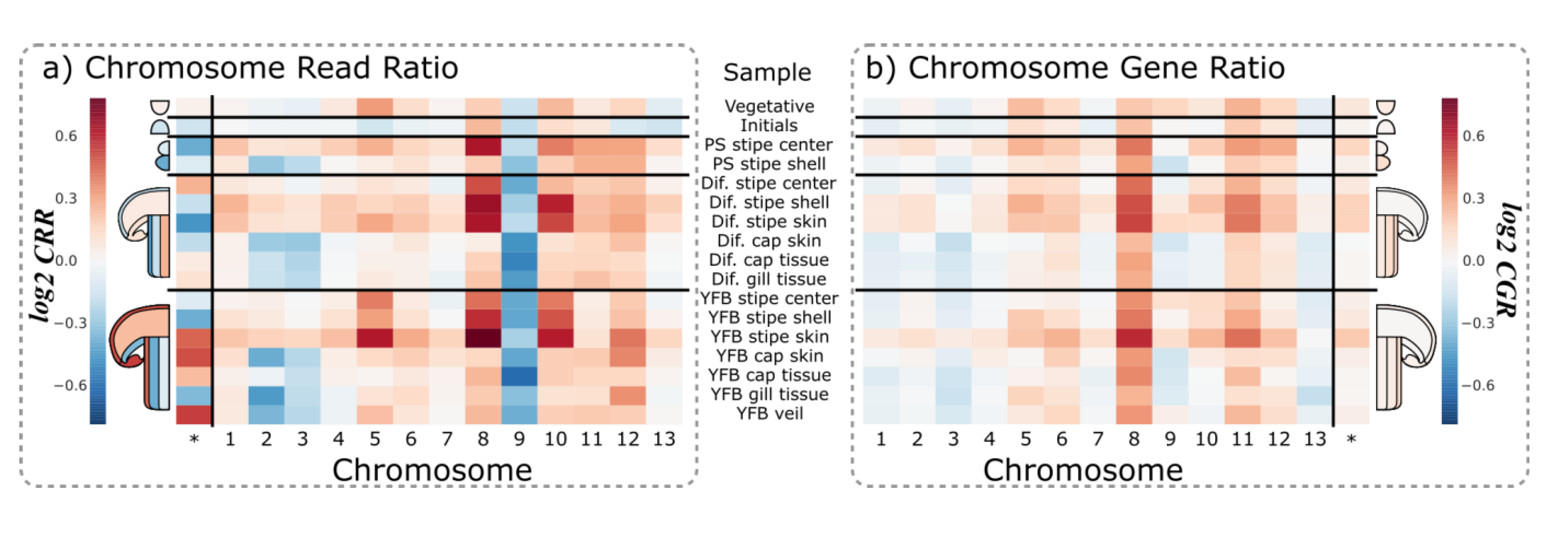
P1 versus P2 expression per chromosome throughout development of the mushroom. A red color indicates a higher P1 activity and a blue color indicates a higher P2 activity. Each row indicates a different developmental stage, and each column represents a different chromosome. The column noted with an asterisk is the ratio at the nuclear type level. **a)**The read count ratios at the chromosome level (CRR, Equation 4). Supplementary Material Note G provides the read count ratios at the chromosome level in the vegetative mycelium dataset. **b)**The Chromosome Gene Ratios (CGR, Equation 6). See Supplementary Material Note H for the gene ratio measures in the vegetative mycelium dataset.

### Gene read ratios reveal a dominant P1 type in mushroom tissue, but not in mycelium

To investigate whether either nuclear type is truly dominant we correct for extremely highly expressed genes (Supplementary Material Note E-F) by limiting their impact on the chromosome and tissue level ratios by using per-gene activity ratios per chromosome (CGR, Equation 6), instead of read ratios. This revealed that, in addition to P1 producing more mRNA than P2, P1 karyolleles were also more frequently higher expressed than their P2 counterpart (Figure 2b). Looking across all tissues and chromosomes, P1 is significantly dominant over P2, i.e. the average of the log-transformed CGR is significantly larger in the P1 nuclear type than the P2 nuclear type, following a t-test in mushroom tissue, with p < 0.01, (see Supplementary Material Note G). Using the Chromosome Gene Ratio has a notable impact on chromosome 9. Although P2 produces most chromosome 9 mRNA (Figure 2a), it is not the case that more P2 karyolleles are more highly expressed than P1 karyolleles.

We do not observe such a dominance of P1 in the mycelium (p > 0.05, with t-test as in mushroom dataset), where neither P1 nor P2 show a dominant mRNA activity (see Supplementary Material Note H).

### A substantial portion of karyolleles are differentially expressed

In each tissue, we determined the set of karyolleles which are statistically significantly differentially expressed between the two nuclear types. Although the dominance of the P1 nuclear type indicates a general trend of higher activity across many genes, some karyollele pairs have a much larger difference pointing towards a functional role. In total, we find 411 genes that are differentially expressed (see Methods) in a mushroom tissue or in vegetative mycelium throughout development (Table 1); 368 genes are differentially expressed in mushroom tissues, and 82 in the vegetative mycelium. The set of differentially expressed genes is enriched in the set of genes with mixed methylation states (Methods, Supplementary Material Notes I).

**Table 1.** Karyolleles differentially expressed between P1 and P2 in mushroom tissue and vegetative mycelium across development. In the first row we indicate the number of differentially expressed genes that are higher expressed in the different nuclear types for the two datasets (columns). The second row gives the total number of differentially expressed genes in the two different datasets. Row three shows the number of differentially expressed genes in a dataset that are not differentially expressed in the other dataset. In the last row, we show the number of differentially expressed genes that overlap between the two datasets.

|  | Mushroom tissue dataset |  | Mycelium tissue dataset |  |
| --- | --- | --- | --- | --- |
|  | P1 up | P2 up | P1 up | P2 up |
| Diff. ex. | 176 | 193 | 30 | 52 |
| Total/dataset | 368 |  | 82 |  |
| Unique/dataset | 329 |  | 43 |  |
| Overlap | 39 (411 total) |  |  |  |

Interestingly, when a karyollele pair is differentially expressed, with only a few exceptions (see Supplementary Material Notes J), it will always be observed to be more highly expressed in the same nuclear type, i.e. if a gene is observed to be more highly expressed in P1 than in P2, than it will never be observed to be more highly expressed in P2 than in P1 in other tissues, and vice versa. The only exceptions to this rule lies in the set of genes that are differentially expressed in both the mushroom dataset and the mycelium dataset.

The set of differentially higher expressed genes between the nuclear types in mushroom and mycelium sets overlap with only 39 genes. In this intersection set, more genes are higher expressed in P2 than in P1. Ten genes had a higher expression in P1, and 24 had a higher expression in P2. Five were more highly expressed in P2 in the mycelium, but switched their origin of primary expression to P1 in the mushroom (see Supplementary Material Notes J). The lack of a substantial overlap of differentially expressed genes between the two nuclear types is indicative of different regulatory processes during the vegetative stage and a mushroom stage.

Although P2 upregulates more differentially expressed genes than P1 does, more genes show a consistently higher expression in P1 than in P2. We identify consistently higher expressed genes that show a higher expression in one nuclear type over the other across all samples (Methods). In the mushroom tissue dataset, we find 1,115 genes that are consistently higher expressed in P1, and 785 genes that are consistently higher expressed in P2. Similarly, in the vegetative mycelium, we find 832 genes that are consistently higher expressed in P1 and 645 that are consistently higher expressed in P2. The two datasets overlap with 470 and 256 genes for P1 and P2, respectively. Interestingly, Of the 90 named genes in *S. commune* (Methods), only *mnp1* is differentially expressed and exhibits different behavior in the mushroom and the vegetative mycelium (see Supplementary Material Note K).

### Co-localized gene clusters are co-regulated

To investigate the level at which genes are regulated, we investigated whether there are regions where the majority of genes were consistently higher expressed in one homokaryon than in the other. We detected many of such regions, given in Table 2 and Figure 3 (Methods, Supplementary Material Note L), hinting towards a sub-chromosomal level of regulation. This is supported by observations in Figure 2, where we see that within one tissue chromosomes are differently regulated, excluding a regulation at the nuclear level. Because we observe that co-regulated gene are co-localized in regions, regulation can also not occur at the chromosome level, because then we would have expected regions of co-regulation of the size of whole chromosomes.

**Figure 3.**
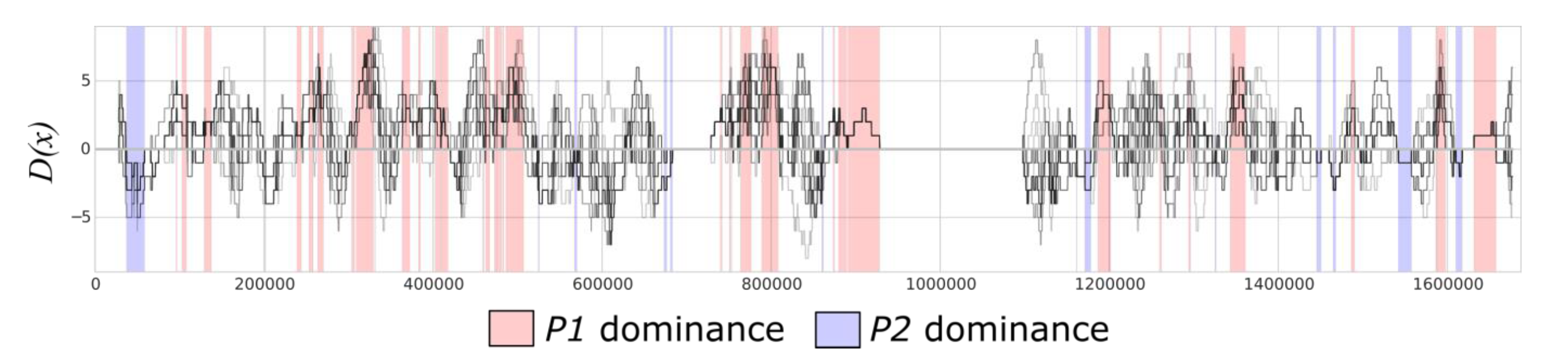
Co-localized genes are often co-regulated. Pictured here are the co-localized and co-regulated gene clusters along chromosome 10 in the mushroom tissue dataset. Along the x-axis is the genomic co-ordinate. For each sample (gray lines), we plot the difference between the number of genes more highly expressed by P1 and the number of genes more highly expressed by P2 (Equation 8, a value of 0 indicates an equal distribution). We also highlight the regions that are consistently upregulated in P1 (red regions) and the number of genes that are consistently upregulated in P2 (blue regions). See Supplementary Material Note J for other chromosomes.

**Table 2.** The number of regions in which the majority of the genes are coregulated (Methods), across the mushroom and mycelium datasets and with the number of genes in these regions. P1 and P2 columns indicate whether the region is consistently higher in for the P1 kayollele or the P2 karyollele, respectiverly. Row Both indicates overlapping regions between the mushroom and vegetative mycelium datasets. Supplementary Material Note L offers detailed expression profiles of these regions.

|  | **P1** |  | **P2** |  |
| --- | --- | --- | --- | --- |
| **Dataset** | **#Regions** | **#Genes** | **#Regions** | **#Genes** |
| **Mushroom** | 207 | 741 | 73 | 233 |
| **Vegetative Mycelium** | 414 | 1955 | 43 | 140 |
| **Both** | 151 | 484 | 7 | 17 |

Co-regulated regions are more frequently upregulated for the P1 karyollele than for the P2 karyolleles. This observation is in agreement with the observed P1 nuclear type dominance. We observe relatively little overlap between the Mushroom and Vegetative Mycelium datasets (Table 2), indicative of different regulatory programs between the vegetative mycelium and mushroom tissue cells.

### Broad range of functionality affected by karyollele specific expression throughout development

Next, we set out to examine the functional annotations of the differentially expressed karyollele pairs, considering the following categories: (i) transcription factors, (ii) metabolic genes, (iii) secondary metabolism genes, (iv) cytochrome P450 genes, (v) carbohydrate active enzymes (cazymes) and (vi) secreted proteins. These categories, with the exception of secondary metabolite genes, are all enriched in the set of differentiable genes (p < 0.05 by a chi-squared approximation to the fisher’s exact test with FDR correction).

Figure 4 show the division of the 411 differentially expressed genes across the functional categories in all the different samples. None of the differentially expressed genes were transcription factors. For the other functional categories, we saw a more or less equal amount of up-regulated karyolleles in P1 and P2 (Figure 4a) in the mushroom tissues (except the vegetative stage), and a more skewed distribution of activity in the mycelium dataset (and the vegetative stage of the mushroom dataset). In these cases, P2 had more differentially expressed genes in these functional categories (Figure 4b).

**Figure 4.**
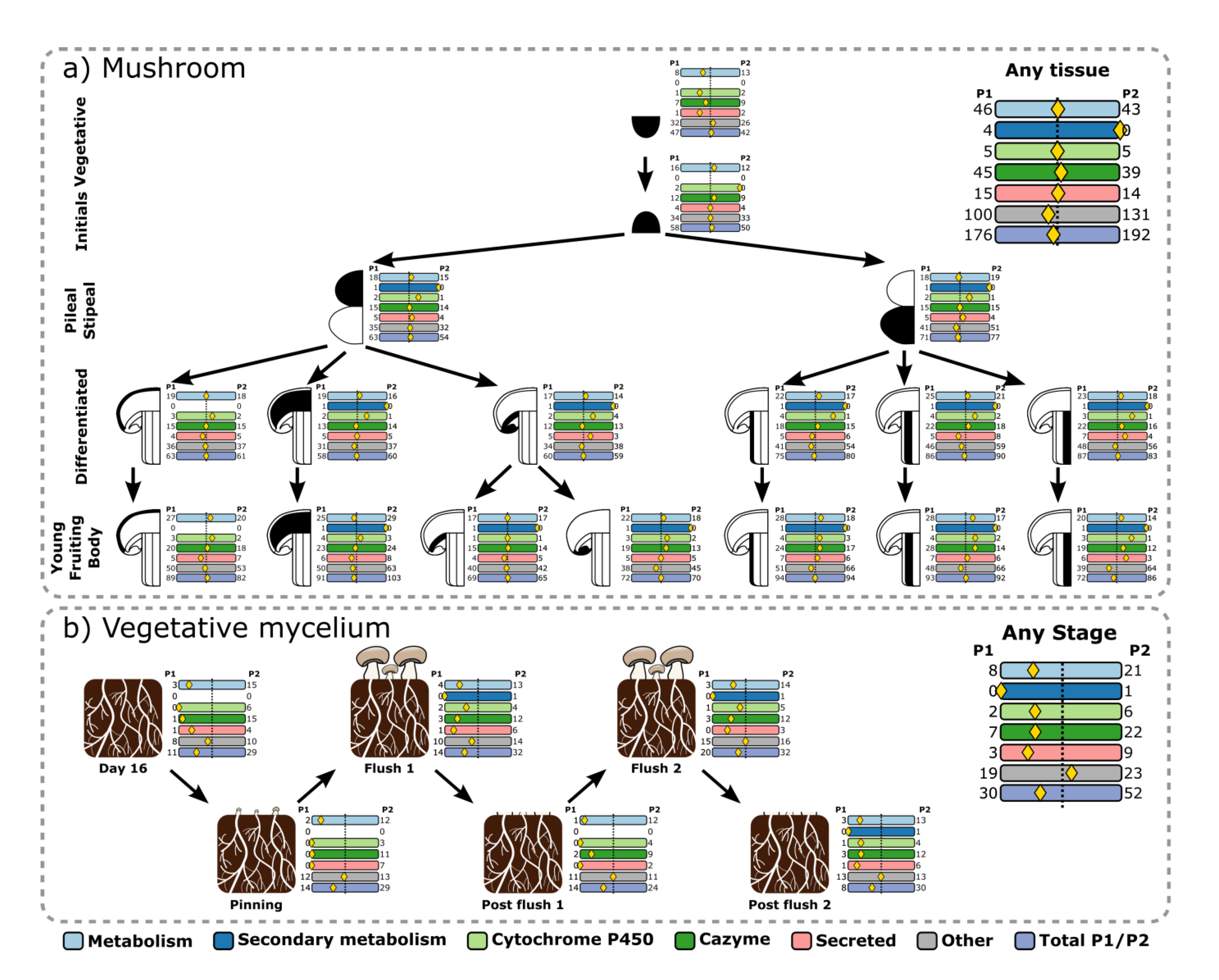
Differential regulation of functional groups through mushroom development. The development of different tissues is illustrated as a tree. We investigate metabolic genes (light blue), secondary metabolic genes (dark blue), cytochrome P450 genes (light green), carbohydrate active enzymes (dark green), secreted protein genes (light red), and all others not fitting into any of the previous groups (grey). At each developmental stage, we observe how many genes of each group are differentially upregulated in P1 (left) and in P2 (right). The yellow diamond indicates the ratio of these counts. **a)** For the mushroom dataset. **b)** For the vegetative mycelium dataset. We see that the groups are more or less equally distributed between P1 and P2 (the yellow diamond is centered), with the exception of the vegetative stage (the root node of Figure 4A), and the vegetative mycelium dataset.

The P2 type had a higher expression of significantly more karyolleles than P1 in mycelium (see Supplementary Material Notes M). In the mycelium, P2 had an enriched expression of cytochrome P450 genes, secondary metabolite genes, and cazymes (p < 0.05, with an FDR corrected chi-squared approximation to the fisher’s exact test). Furthermore, cazymes and metabolic genes in mycelium were more likely to be more highly expressed in P2 (p < 0.05, with an FDR corrected binomial test).

Nineteen of the 39 previously identified differentially expressed genes that are shared between the mycelium and mushroom datasets had the following functional annotations: 14 were annotated as metabolic genes, 14 as cazymes, five as secreted proteins, and two as cytochrome P450s (some genes have multiple annotations). Additionally, five of these 39 overlapping genes have different domain annotations, indicating different functional properties between the P1 and P2 karyolleles.

To further elucidate the functional impact of the 411 differentially expressed genes, we mapped them onto the KEGG pathway database. Sixteen of the genes that are differentially expressed in mushroom tissue or vegetative mycelium samples are found in 20 pathways. Interestingly, three differentially expressed genes are found in the Aminoacyl-tRNA biosynthesis (M00359) pathway (Supplementary Material Notes N). Two genes belong to valine and methionine tRNAs pathways and were upregulated in P1. One gene in the pathway producing aspartamine tRNAs pathway was upregulated in P2. Together, this suggests that P1 is able to produce more valine and methionine tRNAs than P2.

Next we studied whether differential expression of a karyollele also resulted in the production of a functionally different protein due to sequence differences between the karyolleles. 216 of the 5,090 distinguishable karyolleles had sequence differences that led to an alternative protein domain annotation, and 36 of these 216 have alternative domain annotations. 36 of these 216 karyollele pairs are differentially expressed between P1 and P2 (see Supplementary Material Notes O).

## Discussion

Differently from most eukaryotes, nuclei remain side by side during most of the life cycle of basidiomycete fungi. Whether each nucleus is contributing equally to the phenotype and, if not, how this is regulated is largely unknown. In an attempt to understand this, we studied the expression of alleles in both constituent nuclei (P1 and P2) of the button mushroom cultivar Sylvan 15. From the observed average gene expression, we conclude that the expression of nuclear type P1 of the *Agaricus bisporus Sylvan A15 strain* is dominant over nuclear type P2. Remarkably, this dominance is present across all developmental stages in the heterokaryon. We can link this phenomenon to the human case, where in fibroblasts ^29^, it has been shown that individual cells preferentially express one allele over the other, which is not evident over a collection of many cells. Whereas in a diploid genome the cell must rely on heterochromatin DNA packing and RNAi regulatory pathways^30^, heterokaryotic cells could instead control the energy usage of a specific nuclear type.

In the mushroom tissue dataset, the number of up-regulated karyolleles in P1 is approximately equal to those in P2, but in the vegetative mycelium dataset, P2 has more up-regulated karyolleles relative to P1. The contrast between a dominant P1, yet more differentially over–expressed genes in P2 in mushroom tissue is paradoxical. However, there are many genes that show a consistently higher expression in either P1 or P2, with more genes showing a consistently higher expression for P1. Is it possible that the P1 homokaryon is responsible for the basal mRNA production, while P2 plays a more reactive regulatory role? Mechanisms for this kind of regulation are not known. In plants, sub-genome dominance may be linked to methylation of transposable elements^24^. Might it be possible that something similar happens in *A. bisporus?*

Although an imbalance in the number of nuclei could very well explain the dominance of P1, we have shown that genes that are consistently higher expressed in one of the karyolleles do co-localize in sub-chromosomal regions. If there were more P1 nuclei than P2 nuclei, we would have expected a general higher expression of genes of one nuclear type across all chromosomes, which we do not observe.

For many differentially expressed genes, the protein sequence differences between the two karyolleles in the two nuclear types encode for different protein domains. This suggests a functional impact of karyollele specific expression. We also observe a broad range of functionality being differentially expressed between the P1 and the P2 nuclear types. For example, the P2 upregulation of cazymes and metabolic genes in P2 in compost highlight the importance of the P2 homokaryon in development. H97, one of the homokaryons in the cultivar Horst U1, from which Sylvan A15 is derived, displays stronger vegetative growth characteristics than its counterpart H39^1^. This metabolic strength may be passed down from the H97 homokaryon to the Sylvan A15 P2 homokaryon, and the differentially expressed karyolleles may in part be responsible for this. *mnp1,* for example, is an important gene for growth on compost and P2 has indeed inherited the relevant chromosome 2 from H97 (Sonnenberg et al., 2016). Such characteristics are relevant for breeding strategies.

Surprisingly, *mnp1* is expressed and even up-regulated in the mushroom tissues. *mnp1* is known to be involved in lignin degradation, which occurs in the vegetative mycelium ^2,28^. In compost, the abundance decreases dramatically throughout development (Supplementary Material Note K). Therefore, the abundance of *mnp1* in the stipe of the fruiting body is unexpected, although it has been shown that proteins produced in the mycelium can find their way into the mushroom31.

However, it does not explain the fact that the P1 karyollele exists in higher abundance in the mushroom tissues, while the P2 karyollele is higher expressed in the vegetative mycelium. Transport of the P2 karyollele from the vegetative mycelium into the mushroom conflicts with the abundances of the P1 karyollele observed in the mushroom tissues.

A significant proportion of differentially methylated karyolleles were also differentially expressed, most differentially expressed genes are not observed to be methylated. The overlap we observe between methylated genes and differentially expressed genes in different developmental stages explain an effect in the mushroom tissue. However, we cannot link the methylation to a preference of nuclear type. For example, the five differentially expressed genes between compost and mushroom that change their nuclear dominance are not methylated. Although, methylation seems to play a role in the differential use of nuclear type for mRNA production, it only explains 10% of the observed differential expression. This may be due to a limitation of our methylation dataset, (which only comprises vegetative growth), but it may also hint towards other regulatory mechanisms.

In addition to methylation, we also observe co-localization of co-expressed genes. This may be indicative of a difference in genome organization, whereby the DNA is less accessible in certain regions in P1 than in P2 through different levels of chromatin compaction. It has been shown that gene expression is strongly linked to DNA availability, and further, that such chromatin organization is heritable^32^.

The sequences of a pair of karyolleles need to be sufficiently different for our algorithm to be able to uniquely assign reads to each karyollele. These sequence differences between nuclear types may have an effect on various regulatory mechanisms of transcription, such as transcription factor binding efficiencies, transcription efficiency, differences in mRNA stability, or differences in epigenetic factors. Future research might shed light on whether these differences are related to observed differential karyollele expression.

Causative mechanisms of karyollele specific expression can further be elucidated by population studies across multiple spore isolates. Sylvan A15 is derived as a heterokaryotic single spore isolate from Horst U1. In such heterokaryons, non-sister nuclei are paired in one spore. Combined with the restriction of recombination to chromosome ends, such heterokaryons are genetically very similar to the parent and differ only in the distribution of parental type chromosomes over both nuclei. Karyollele expression could thus be studied in different heterokaryotic single spore isolates having different distributions of otherwise very similar chromosomes over both nuclei. If the expression patterns are consistent with nuclear chromosome organization across different single spore isolates, it will suggest that expression of specific karyolleles can be controlled by selecting isolates where karyolleles lie in the desired nuclei.

## Conclusion

We show that karyolleles, the different copies of a gene separated by nuclear membranes in a heterokaryon, are differentially expressed between the two different nuclear types in the *Agaricus bisporus Sylvan A15 strain.* Each nuclear type contributes varying amounts of mRNA to the cell, and differential expression occurs at the gene level. Despite a dominant P1 type, we see no evidence that would suggest an imbalance in the number of copies P1 and P2 nuclei in any cell type, though it may vary from cell to cell.

Genes with various vital functions are differentially expressed. The P2 homokaryon significantly up-regulates cazymes and metabolic genes, which may indicate a difference in vegetative growth strengths. This corroborates what was observed in the constituent homokaryons of the Horst U1 cultivar from which P1 and P2 are essentially derived.. Manganese peroxidase is one of the differentially expressed genes, and exhibits interesting, previously unknown behavior. The cause of these differential regulations is still not known, but it is possible that epigenetic mechanisms, like methylation, play a role.

The biological gene regulation mechanisms between heterokaryons need to be investigated. Unfortunately, such research is hindered by current mRNA isolation procedures. As mRNA transcripts are secreted from the nuclei and mixed in the cytoplasm of the cell, traditional sequencing methods will be unable to generate a full resolution of both homokaryon expression from full cell isolates. Single nucleus sequencing ^33,34^ would circumnavigate this issue by isolating mRNA from individual nuclei. As we have shown that the two nuclear types exhibit distinguishable regulatory programs, it will be possible to distinguish them based on their expression profiles.

The impact of differential expression between nuclei of heterokaryotic organisms is underappreciated. Heterokaryotic fungi have major impact in clinical and biotechnological applications, and impact our economy and society as animal pathogens such as *Cryptococcus neoformans*^35^, plant pathogens such as *Ustilago maydis^36^,* plant and soil symbionts such as mycorrizal fungi^26^, bioreactors such as *Schizophyllum commune^31^,* and of course the subject of this study, the cultivated, edible mushroom *Agaricus bisporus*^15^. It is known that different homokaryons in these species will produce different phenotypes^2^ which no doubt need to be treated, nourished or utilized differently.

We have demonstrated differential nuclear regulation of a fungal organism and we showed that variation between homokaryons results in functional differences that were previously unknown. With this work, we hope to draw attention to the impact of sequence and regulatory variation in different nuclei on the function and behavior of the cell in order to further our understanding of the role of fungi in our environment.

## Materials and Methods

### RNA-Seq data

We used two RNA-seq datasets from the *Agaricus bisporus (A15)* strain: (1)tissue samples through mushroom development (BioProject: PRJNA309475)^27^, and (2) vegetative mycelium samples taken from compost through mushroom development (BioProject PRJNA275107)^2^. Throughout the text, when we refer to the mushroom tissue, we also refer to all samples in dataset (1), including the first sample, which technically is a sample of the vegetative mycelium. The compost dataset exhibited high amounts of PCR duplicates (Supplementary Material Note P). This can be attributed to the difficulty in isolating RNA from soil. To remedy the biases involved with this, we removed all PCR duplicates using FastUniq ^38^.

### Methylation data

A sample of vegetative stage mycelium of A15 was treated with the EpiTect Bisulphite conversion and cleanup kit and sequenced with the Illumina HiSeq 2000. Raw reads were trimmed using TRIMMOMATIC^39^ and aligned to the A15 P1 genome using Bismark^40^ and bowtie2^41^. Methylated bases were analyzed with Methylkit^42^. Only bases which had a minimum coverage of 10 were retained. For samples with mixed methylation states, we will observe what appear to be incomplete conversions of unmethylated cytosines but in reality represents the mixed methylation states of those bases. Therefore, to include only differentially methylated bases between the two nuclei (i.e. methylated in one homokaryon, but not in the other), we considered only those bases which were measured to be methylated between 40 and 60% of all reads (Supplementary Material Notes I). While 164,290 bases had an indication of methylation signal, 10,325 bases had methylation signals of about 50%, suggestive of differential methylation states. Methylated bases were mapped to genes when between the start and stop codons, or 1000bp up/downstream (Supplementary Material Note Q).

### Homokaryon genome and annotations

The P1 and P2 genomes^15^ were annotated with BRAKER1^43^ using the pooled RNA-seq data described above. In order to prevent chimeric genes (neighboring genes that are erroneously fused into one predicted gene) the following procedure was used. After the first round of gene prediction, predicted introns were identified that were at least 150 bp in size and not supported by RNA-seq reads. The midpoint of these introns were labeled as intergenic regions in the next round of gene prediction using AUGUSTUS 3.0.2^44^ and the parameter set produced in the first round of gene prediction. The SNP density between the genomes was estimated using MUMMER’s^45^ show-snps tool.

### Karyollele pair discovery

The genome annotations were used to produce predicted mRNA sequences for each gene. The genes in the two parental genomes were matched using a reciprocal best BLAST ^46^ hit. Hits which had E-values greater than 10^-100^ were removed. This resulted in a conservative orthology prediction between the two homokaryons that are our set of karyolleles. Karyollele pairs which have a 100% sequence identity were removed, as it would be impossible to identify distinguishing markers for these identical pairs.

### Marker Discovery

For each discovered karyollele pair, we identify markers that uniquely identify each element of the pair. This is done by constructing all possible kmers for each sequence, resulting in two sets per pair. The kmers overlapping in these sets are removed, resulting in distinguishing pairs of markers. Once distinguishing markers have been discovered for all pairs, we remove all non-unique markers. Finally, the set of markers is made non redundant by scanning the position-sorted list of markers from left to right and removing any marker that overlaps with the previous marker. Finally, we ensure that the markers are unique throughout the whole genome by removing markers that are present anywhere else in either genome. In order to guarantee sufficient evidence across the whole gene, we remove karyollele pairs which do not have at least five markers each.

### Marker quantification

We scan all RNA-Seq reads for the detected markers using the Aho-Corasick algorithm^47^. We insert all markers and their reverse complements into an Aho-Corasick tree and count each marker only once for each fragment (a marker may be present twice, if the read mates overlap). We calculate a gene expression score as the average of each marker count for a gene. This results in an expression score *E_h_* for each gene *g* in each sample *t* for each replicate *r*, per homokaryon *h*:
 

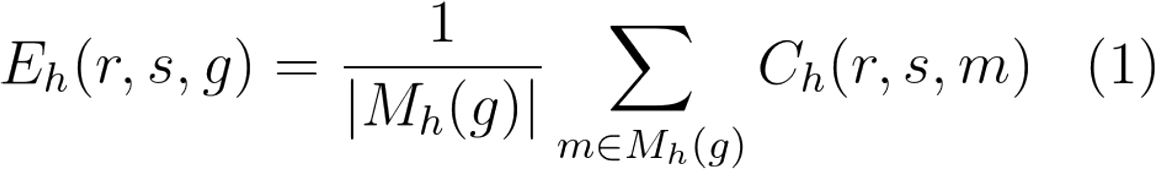

where *M_h_(g) is* the set of markers in a gene *g,* and *C_h_(r,s,m)* is the count for marker m in replicate *r,* sample *s*.

### Differential expression

Using DE-Seq^48^, we perform a differential expression test for each karyollele pair in a tissue, i.e. we test if a gene has a differential expression in P1 or P2. DESeq requires a size factor to be calculated, which normalizes for the library sizes of each sample. Since however, the counts from P1 and from P2 originate from the same sample, these must have the same size factor. Size factors are therefore calculated manually, by counting the total number of reads for each sample, and dividing it by the largest value for any sample (Equation 2).

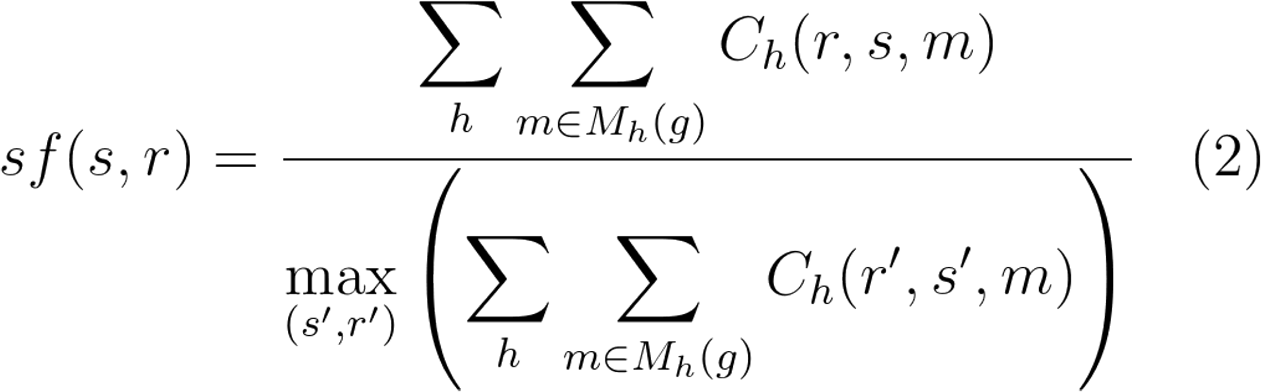

The P1 and P2 counts originating from the same sample will then be assigned the same size factor. The expression counts for each gene in each replicate in each tissue (equation 1) are provided to DE-Seq with the provided size factor (Equation 2). The normalized read counts per gene *D_h_(s,g)* are returned by DE-Seq, together with significance values for each test. We select only differentially expressed genes that have a q-value < 0.05, and a fold change of at least three.

### Read ratio calculation

Using the normalized read counts from DE-Seq ^48^, we calculate the ratio of the number of reads originating from the two homokaryons at the gene (GRR), chromosome (CRR) and nuclear type level (NRR).

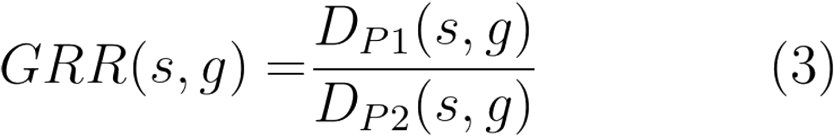

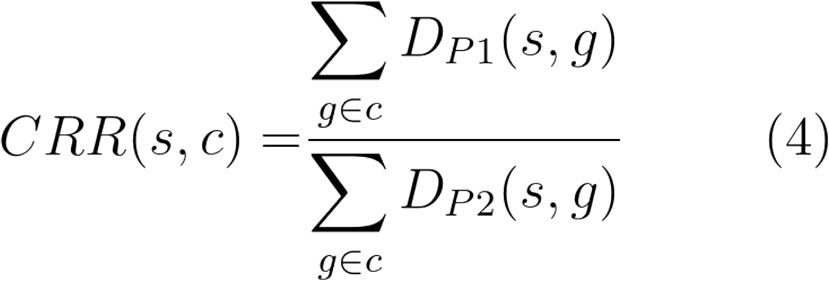

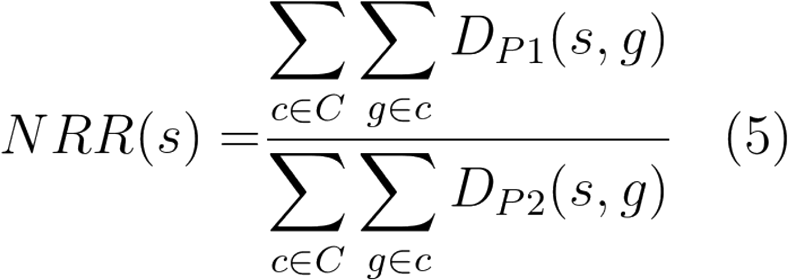

### Gene ratio calculation

Using the normalized read counts from DESeq^48^, we calculate the ratio of the number of reads originating from the two homokarons at the gene level, and use those ratios to calculate the geometric mean of the relative expression activities at the chromosome (CGR, Equation 6) and nuclear type level (NGR, Equation 7). The geometric mean is more suitable than the arithmetic mean for averaging ratios.

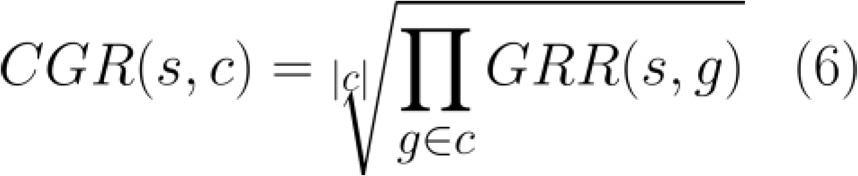

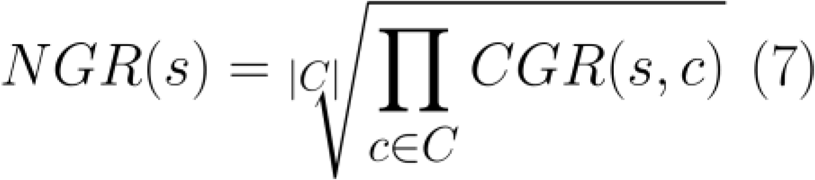

### Identifying consistent genes

For each gene, we observe the relative expression in each sample (Equation 3). We refer to a gene as being consistently expressed if it is more highly expressed in the same nuclear type in each sample. I.e. the GRR is always greater than one, or always less than 1.

### Identifying co-regulated clusters

We slide a window of size 20,001bp (10,000- up and down-stream) across each chromosome. In this window, we count the number of genes that are more highly expressed by P1 and by P2, and calculate the difference per sample. I.e.

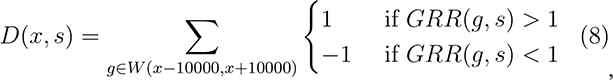

Where *W(x,y)* is the set of genes between genomic location x and y, and s is a sample. This difference is shown in Figure 3. Next, we identify regions where each sample in the dataset shows consistent regulation. That is to say, in these regions, 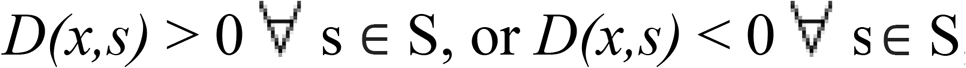, where S is the set of all samples. These regions contain co-localized genes that are co–regulated across all samples.

## Functional predictions

### PFAM

Conserved protein domains were predicted using PFAM version 27^49,50^.

### Transcription factor definitions

Predicted proteins with a known transcription factor-related (DNA-binding) domain (based on the PFAM annotations) were considered to be transcription factors.

### Carbohydrate-active enzymes prediction

Using the Cazymes Analysis Toolkit (CAT) ^51^, we predicted carbohydrate-active enzymes based on the original gene definitions. If a gene’s protein sequence was predicted to be a cazyme by either the sequence-based annotation method or the PFAM-based annotation method then we considered it a cazyme.

### Secreted Proteins prediction

We used the same procedure as ^52^ to predict secreted proteins. Briefly, genes with SignalP ^53^ signal peptides, or a TargetP ^54^ Loc=S were kept. The remaining genes were further filtered with TMHMM ^55^, keeping only genes with zero or one transmembrane domains. Finally, genes were filtered using Wolf PSort ^56^ to select genes with a Wolf PSort extracellular score greater than 17.

### Metabolic and Cytochrome P450 gene groups

Genes with the GO annotation “metabolic process” (annotation ID: GO:0008152) were called as metabolism genes. Genes with the PFAM annotation PF00067 were used as Cytochrome P450 genes.

### KEGG

KEGG annotations were made with the KAAS KEGG^57^ annotation pipeline, using genes from all available fungi, with the exception of leotiomycetes, Dothideomycetes, and Microsporidians, due to the limitation of the number of species (Selected organisms by ID: cne, cgi, ppl, mpr, scm, uma, mgl, sce, ago, kla, vpo, zro, cgr, ncs, tpf, ppa, dha, pic, pgu, lel, cal, yli, clu, ncr, mgr, fgr, nhe, maw, ani, afm, aor, ang, nfi, pcs, cim, cpw, pbl, ure, spo, tml). The GHOSTX and BBH options were selected. Predictions were made individually for both the P1 and P2 genomes, using the translated protein sequences.

### Named genes

Named genes for *Agaricus bisporus* version 2 were downloaded from the JGI DOE Genome Portal (http://genome.jgi.doe.gov/pages/search-for-genes.jsf?organism=Agabi_varbisH972) by searching for genes with ‘Name’ in the ‘user annotations’ attribute. Gene names were transferred from *A. bisporus* v. 2 using reciprocal best blast hit to P1 and P2, and then selecting the best match (in the single case of an ambiguity). See Supplementary Material Note R.

### Software and code availability

Marker discovery and abundance calculations was done in Scala, while downstream analysis was performed in python using the ibidas data query and manipulation suite^58^. All source code, together with a small artificial example dataset is available at: https://github.com/thiesgehrmann/Homokaryon-Expression

### Data Availability

The RNA-Seq data was previously generated and can be found at bioprojects PRJNA309475 and PRJNA275107. The bisulphite sequencing data can be accessed at SAMN06284058.

### Supplementary information

Together with this manuscript, we provide a file of Supplementary Notes, and Supplementary Tables 1-4 to support our findings.

## Acknowledgements

The authors would like to thank Brian Lavrijssen for providing the bisulphite sequencing data to determine the differential methylation states. The sequence and annotation data of *A. bisporus* H97 version 2 were produced by the US Department of Energy Joint Genome Institute http://www.jgi.doe.gov/ in collaboration with the user community. This research is supported by the Dutch Technology Foundation STW, which is part of the Netherlands Organisation for Scientific Research (NWO), and which is partly funded by the Ministry of Economic Affairs.

## Author contributions

TG, HABW, MJTR and TA wrote the manuscript. JFP performed the experiments. TG, HABW, MJTR and TA designed the analyses. RAO created the gene and functional annotations. TG performed the analyses. All authors aided in biological interpretation of the results. All authors reviewed the manuscript.

